# *In vitro* persistence level reflects *in vivo* antibiotic survival of natural *Pseudomonas aeruginosa* isolates in a murine lung infection model

**DOI:** 10.1101/2022.10.21.513228

**Authors:** Laure Verstraete, Juliana Aizawa, Matthias Govaerts, Linda De Vooght, Jan Michiels, Bram Van den Bergh, Paul Cos

**Author notes:** These authors contributed equally to the work as first authors (Laure Verstraete, Juliana Aizawa). These authors contributed equally to the work as senior authors (Jan Michiels, Bram Van den Bergh, Paul Cos).

## Abstract

Nowadays, clinicians are more and more confronted with the limitations of antibiotics to completely cure bacterial infections in patients. It has long been assumed that only antibiotic resistance plays a pivotal role in this. Indeed, the worldwide emergence of antibiotic resistance is considered as one of the major health threats of the 21^st^ century. However, the presence of persister cells also has a significant influence on treatment outcomes. These antibiotic-tolerant cells are present in every bacterial population and are the result of the phenotypic switching of normal, antibiotic-sensitive cells. Persister cells complicate current antibiotic therapies and contribute to the development of resistance. In the past, extensive research has been performed to investigate persistence in laboratory settings, however, antibiotic tolerance in conditions that mimic the clinical setting is still poorly understood. In this study, we have optimized a mouse model for lung infections of the opportunistic pathogen *Pseudomonas aeruginosa*. In this model, mice are intratracheally infected with *P. aeruginosa* embedded in seaweed alginate beads and subsequently treated with tobramycin via nasal droplets. A strain panel of 18 *P. aeruginosa* isolates originating from environmental, human and animal clinical sources was selected to assess survival in the animal model. These survival levels were positively correlated with the survival levels determined via time-kill assays which is a common method to study persistence in the laboratory. We showed that both survival levels are comparable and thus that the classical persister assays are indicative for antibiotic tolerance in a clinical setting. The optimized animal model also allows us to test potential antipersister molecules and study persistence.

**Importance:** The importance of targeting persister cells in antibiotic therapies becomes more evident as these antibiotic-tolerant cells underlie relapsing infections and resistance development. Here, we studied persistence in a clinically relevant pathogen, *Pseudomonas aeruginosa*. It is one of the six ESKAPE pathogens (*Enterococcus faecium, Staphylococcus aureus, Klebsiella pneumoniae, Acinetobacter baumannii, P. aeruginosa, Enterobacter* spp.) that are considered as a major health threat. *P. aeruginosa* is mostly known for causing chronic lung infections in cystic fibrosis patients. We mimicked these lung infections in a mouse model to study persistence in more clinical conditions. We showed that the survival levels of natural *P. aeruginosa* isolates in this model are positively correlated with the survival levels measured in classical persistence assays. These results not only validate the use of our current techniques to study persistence, but also open opportunities to study new persistence mechanisms or evaluate new antipersister compounds *in vivo*.

## Introduction

*Pseudomonas aeruginosa* is a Gram-negative, opportunistic pathogen that is commonly associated with chronic wound infections and airway infections in cystic fibrosis (CF) patients [1, 2]. The intrinsic resistance of *P. aeruginosa* and its ability to acquire additional resistance mechanisms make *P. aeruginosa* infections very challenging to treat [3]. It is considered as a priority pathogen by the World Health Organization for which new treatments are urgently needed and drug research should be prioritized [4]. A recent analysis on the impact of antimicrobial resistance worldwide estimates that in 2019 more than 300,000 deaths are associated with resistant *P. aeruginosa* infections [5]. Not only resistance, but also the presence of a subset antibiotic-tolerant cells may explain the difficulty to eradicate *P. aeruginosa* infections [6, 7]. Indeed, when challenged with a high dose of bactericidal antibiotics, a small fraction of persister cells is able to survive even in the absence of antibiotic resistance. Persister cells are phenotypic variants of normal antibiotic-sensitive cells that have transiently switched to a non-growing, antibiotic-tolerant state. In absence of antibiotics, these cells give rise to a new bacterial population that is equally susceptible to antibiotics as the original population [8, 9]. Due to its small fraction, persisters are often neglected or overlooked but are nonetheless of great clinical concern as they are inherently present in all bacterial populations and underlie antibiotic therapy failure. Persisters are especially relevant in biofilms and intracellular infections, or in immunocompromised patients where the immune system is not able to eliminate all cells [10, 11]. More recently it has been shown that persisters catalyze resistance development in several ways. Persisters form a pool of viable cells from which resistant mutants can emerge via *de novo* mutations [12, 13] or horizontal gene transfer [14]. Moreover, higher persistence levels are associated with higher mutation rates conferring resistance in natural isolates and mutated lab strains of *Escherichia coli* [15]. A positive correlation between resistance and persistence levels has also been observed in natural isolates of *P. aeruginosa* [16], suggesting a complementary link between the two survival strategies. Thus, targeting persister cells not only kills antibiotic-tolerant cells, but also impedes resistance development. As its clinical importance is slowly being acknowledged, recent years have seen an increasing interest in persistence research. Most studies have focused on the mechanisms of persister formation [17] and awakening [18–20] in *E. coli* but the complete picture is still not delineated in full. Also, knowledge on the physiology of persister cells of major pathogens such as *P. aeruginosa* remains fragmentary. Some of the already-known persister mechanisms are condition-dependent [8] and have mainly been observed in lab conditions which raises the question whether the same mechanisms are relevant in real-life infections. Only few studies could extrapolate *in vitro* findings on persistence to an *in vivo* setting [21–23], while others obtained mixed [24] or contradictory [25, 26] results. Current persistence models have mainly focused on lung infections caused by *Mycobacterium* species [26–29] and intracellular infections by *Salmonella* [14, 22, 25, 30–34]. In the past, *P. aeruginosa* persisters were studied *in vivo* using an intraperitoneal and subcutaneous biofilm infection model in mice [21]. However, *P. aeruginosa* is most lethal when infecting the lungs [35] and a lung infection model to study antibiotic tolerance is currently lacking. Moreover, it is unknown whether the number of persisters quantified via standardized persister assays in the lab resembles antibiotic tolerance *in vivo*. To fill this void, we optimized a murine lung infection model with seaweed alginate beads to study antibiotic tolerance. Additionally, we assessed a diverse set of natural *P. aeruginosa* isolates in classical *in vitro* persister assays and compared the results to their survival in the murine lung model. We found that the survival level is highly variable among the strains both *in vitro* and *in vivo*. More importantly, the survival in the murine model positively correlates with the survival measured in laboratory settings. While some deviations between *in vitro* and *in vivo* findings were found, our results show that *in vitro* set-ups to study antibiotic tolerance are appropriate. Moreover, in future studies, the optimized infection model could be used to test new antipersister compounds to combat *P. aeruginosa* infections or to study tolerance mechanisms.

## Results

### Optimization of the murine infection model

Mammalian species, and more specially rodents, are often the preferred animal model to study the *in vivo* pathogenesis of lung infections in a more controlled setting. Mice have been widely used in pulmonary research because of their small size and high reproducibility, the availability of inbred or transgenic strains and the full characterization of the mouse genetics and immunology [36, 37]. To overcome the intrinsic tolerance of mice to human pathogens, the host immune system could be suppressed or physically separated from the bacteria. In this study, bacterial clearance by the murine immune system is prevented by infecting the mice with *P. aeruginosa* embedded in seaweed alginate beads (Figure 1A). This infection model was first developed in rats with bacteria embedded in agar beads [38], adapted for mice [39] and further optimized with seaweed alginate beads [40] and in BALB/c mice [41, 42]. The use of seaweed alginate beads mimics the later stages of chronic lung infections where mucoid *P. aeruginosa* produces alginate which is an important constituent of the extracellular matrix of biofilms [40, 43]. Moreover, a study with agarose beads showed that the bacteria can still migrate from the beads and grow slowly, which is similar to the phenotype in biofilm [44]. In our study, the seaweed alginate solution was directly installed in the lower respiratory tract of the mice via the oral route which prevents animal injury and ensures the direct deposition of the inoculum causing less variation between individual mice [45]. After infection, the mice were treated with high doses of tobramycin, an aminoglycoside antibiotic that is widely used for the treatment of *P. aeruginosa* infections. In patients, tobramycin is administered via inhalation therapy in which the product is directly delivered in the airways [46, 47]. Here, this antibiotic therapy was mimicked by the active inhalation of tobramycin via nasal droplets. Overall, these modifications result in a mouse model that mimics pulmonary infections in patients.

**Figure 1:**
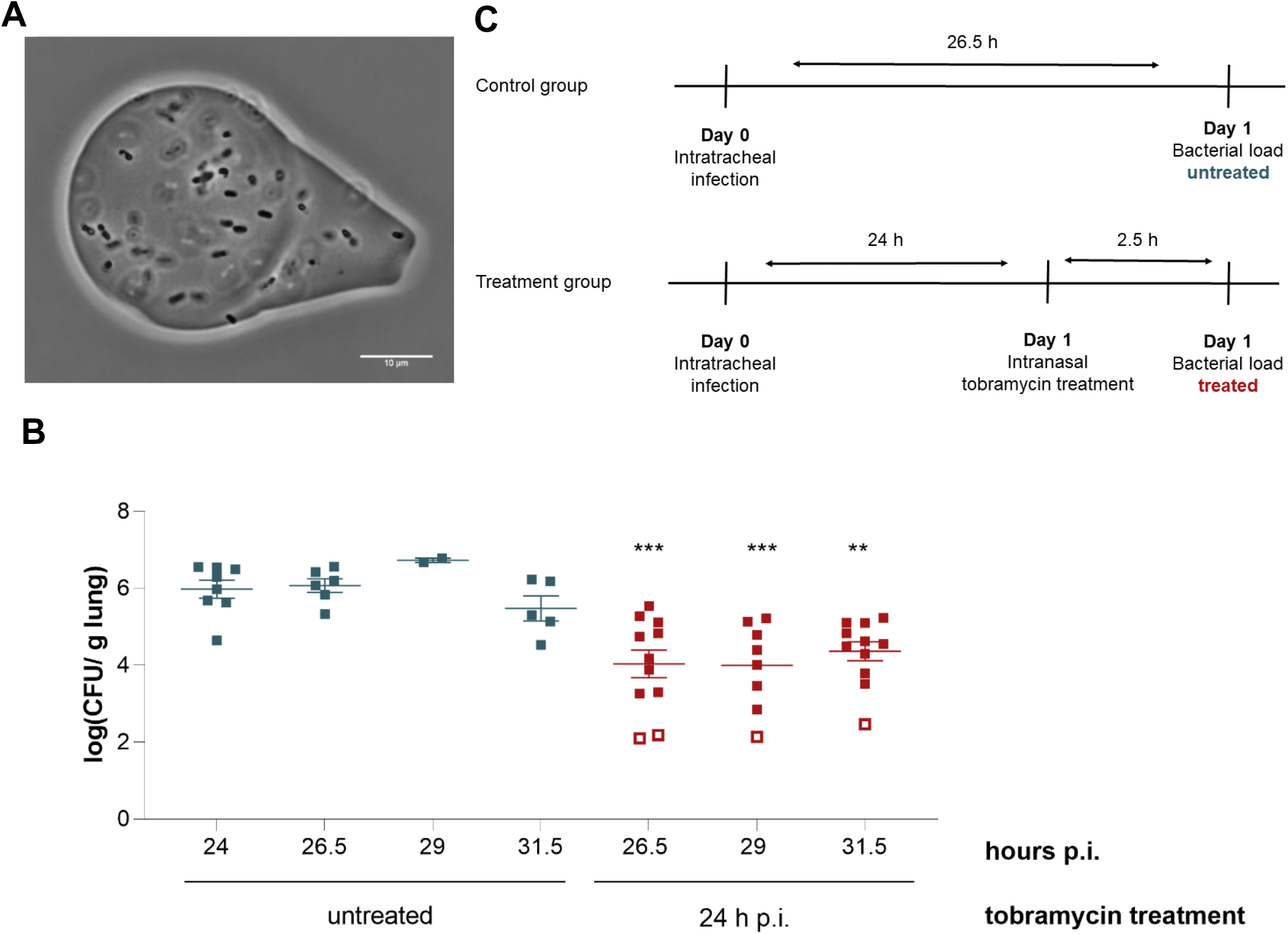
The lung infection model with P. aeruginosa embedded in seaweed alginate beads can be used to study antibiotic tolerance in vivo. (A) Microscopic image of seaweed alginate beads containing P. aeruginosa cells. (B) Pilot experiment with PA14. The bacterial load in the lungs of untreated mice remains constant over time up to 31.5 h p.i. Upon tobramycin treatment, the number of cells in the murine lungs decreases significantly compared to the untreated mice at the same time point. Statistical differences between the bacterial load of untreated mice at 24 h p.i. and treated mice were determined with one-way ANOVA with a Dunnett’s post hoc test for multiple comparisons (**, P < 0.01; ***, P < 0.001). Each symbol represents one mouse and the error bars show the standard error of the mean. Open squares indicate repeats below the detection limit of which half of the detection limit divided by the lung weight is shown. (C) Optimized treatment scheme to determine the survival of antibiotic-tolerant cells in mice.

A first set of conditions were chosen based on the mouse model infected with the lab strain PA14. In a pilot experiment, BALB/c mice were intratracheally infected with several doses of PA14 embedded in seaweed alginate beads to study the infection course. Survival of the mice was measured 48 h post infection (p.i.) and the bacterial load in the lungs was determined by plating. This dose-response experiment demonstrated that a challenge dose of 5×10^5^ CFU/mouse caused a stable infection and resulted in a bacterial titer of around 5×10^6^ CFU/g lung after 48 h. Higher doses resulted in a high mortality (50%-100%), while with lower challenge doses all bacteria were cleared. For consistency, the same dose of 5×10^5^ CFU/mouse was used for all subsequent infections, including the ones with natural *P. aeruginosa* isolates. In the next step, the killing kinetics of bacteria in the lungs upon tobramycin treatment was characterized. A tobramycin dose of 120 mg/kg body weight, the highest tolerable dose, was applied after 24 h of infection with PA14-alginate beads. This tobramycin concentration, equivalent to 60 mg/ml, is at least 100,000-fold higher than the MIC of PA14 (~0.25 μg/ml) which should ensure that antibiotic-sensitive cells are effectively killed and antibiotic-tolerant cells survive. After antibiotic administration, the bacterial load in the lung was measured every 2.5 h (Figure 1B). After 2.5 h, the bacterial load of treated mice decreased 2 log compared to untreated mice. The killing rate slowed down after 2.5 h as the number of CFU in the lung did not further decrease over time. To check whether this CFU reduction is mainly caused by antibiotic exposure and not due to reduced survival in the lungs, the number of bacteria in the lungs of untreated mice was also measured. Indeed, the number of viable cells in untreated samples remained constant over the same period indicating that the applied tobramycin dose effectively kills bacterial cells. The observed killing pattern upon treatment with high doses of antibiotics is consistent with biphasic killing kinetics which is typically associated with antibiotic persistence (Figure S1) [9]. There was no dissemination of bacteria to the liver or spleen observed (data not shown). In subsequent experiments with other *P. aeruginosa* strains, only one euthanasia time point was chosen to reduce animal usage. Since no differences in CFU were observed between the early and late time points, it was decided to treat the mice for 2.5 h and thus kill both untreated and treated mice 26.5 h p.i. (Figure 1C). An agar well diffusion assay demonstrated that the tobramycin concentration remained well above the MIC of PA14 throughout the treatment. The measured concentration of the homogenized lungs, sampled 2.5 h after treatment, was 8 (± 3) μg/g lung. These optimized conditions, i.e. challenge dose, tobramycin concentration and treatment duration, were further used to determine the survival of natural isolates in mice.

### *In vitro* survival of natural *P. aeruginosa* isolates

In a first series of tests, we selected a panel of natural *P. aeruginosa* isolates based on our *in vitro* findings that could later be used for testing in the animal model. In total, 18 isolates were chosen based on three main criteria: MIC levels, strain origin and *in vitro* persistence levels. To exclude antibiotic resistance as a confounding factor in bacterial survival, all strains were selected to be equally sensitive to tobramycin. The MIC of the selected isolates ranges between 0.125 and 0.5 μg/ml (Table 1). To capture the diversity within the *P. aeruginosa* species, these isolates were derived from diverse sources (Table 1). The final strain selection, therefore, includes human clinical strains (isolated from CF patients and wound infections), animal strains (isolated from dogs, kangaroos, parrots and cats) and environmental strains (isolated from water sources and plants). The constructed phylogenetic tree of the isolates showed that strains from the same origin are not phylogenetically related as they do not appear in the same clades, demonstrating the diversity of the strain collection (Figure 2). Finally, the persistence levels vary across the selected isolates which increases the statistical power of the comparison between the *in vitro* and *in vivo* survival. Time-kill experiments were performed to determine the persistence level of each *P. aeruginosa* isolate (Figure 3). In these experiments, cultures were exposed to high doses of tobramycin (100x MIC) for several hours. To obtain a quantitative measure for persistence, the persister fractions were obtained from the estimated parameters of the biphasic models fitted to the time-kill data (Table S1). The persister fractions varied largely among the selected strains (p < 0.0001), ranging from 3×10^−7^ to 0.09, and were not phylogenetic correlated (Figure 2). Human strains have significantly higher survival levels compared to animal or environmental strains (p < 0.001). The strains could be categorized into three groups based on their survival fraction (Table 1). Strains were ordered from the lowest to the highest survival fraction. The first and last six strains are considered as strains with a low and high survival phenotype, respectively. The other six strains with an intermediate survival fraction belong to the intermediate group. The average persister fraction between these three survival groups is significantly different (p < 0.0001) (Figure S2A). In conclusion, a panel of diverse *P. aeruginosa* strains with low resistance levels was assembled. Moreover, a wide diversity in persistence levels between the strains was considered which makes it suitable for comparison with survival *in vivo*.

**Figure 2:**
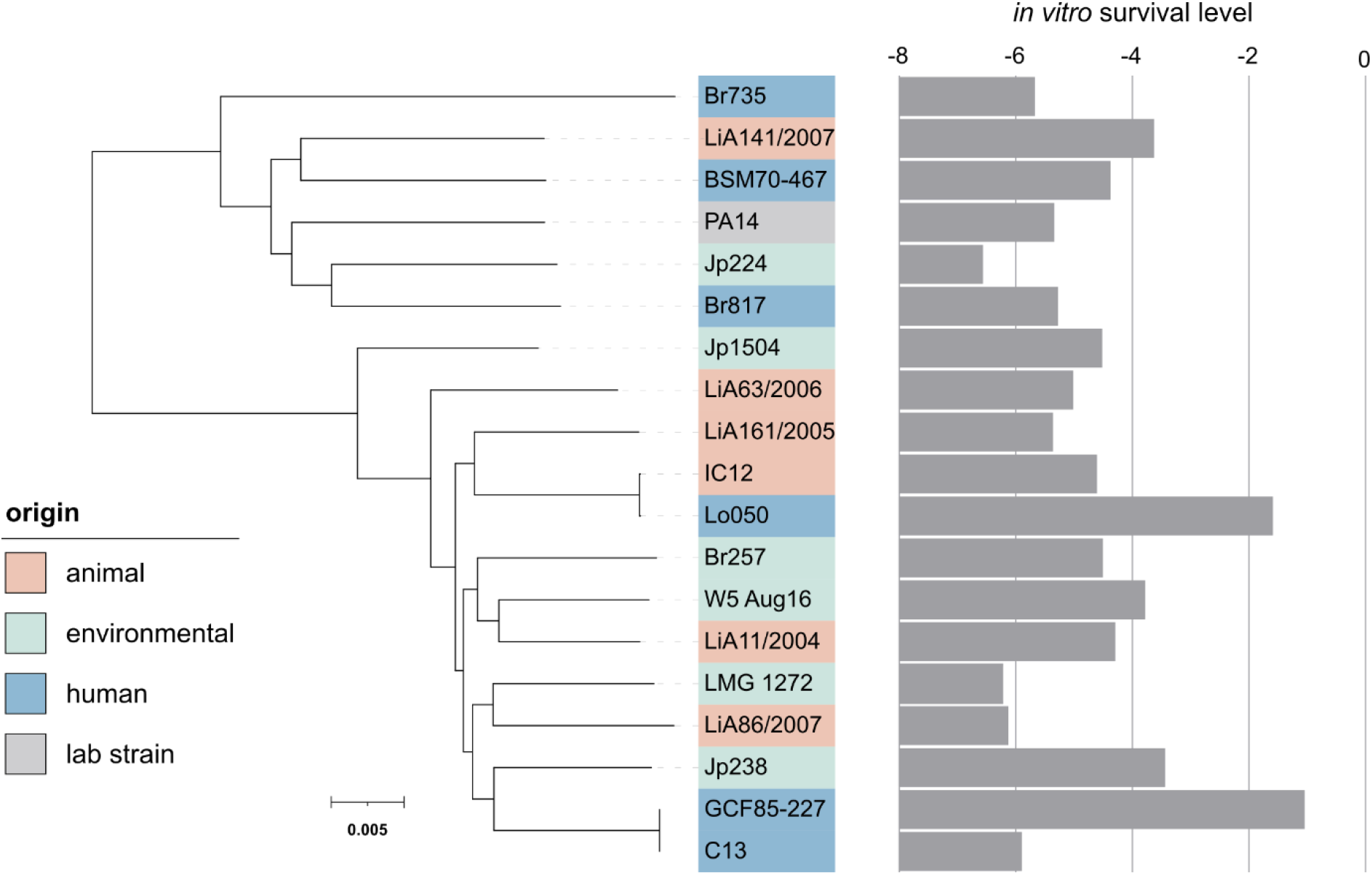
Phylogenetic tree based on the core genome of the P. aeruginosa strains used in this study. All isolates are colored according to their origin and the bar plot shows the in vitro survival. PA7 is used as an outgroup to root the tree and is not shown. The scale bar indicates the number of substitutions per site.

**Figure 3:**
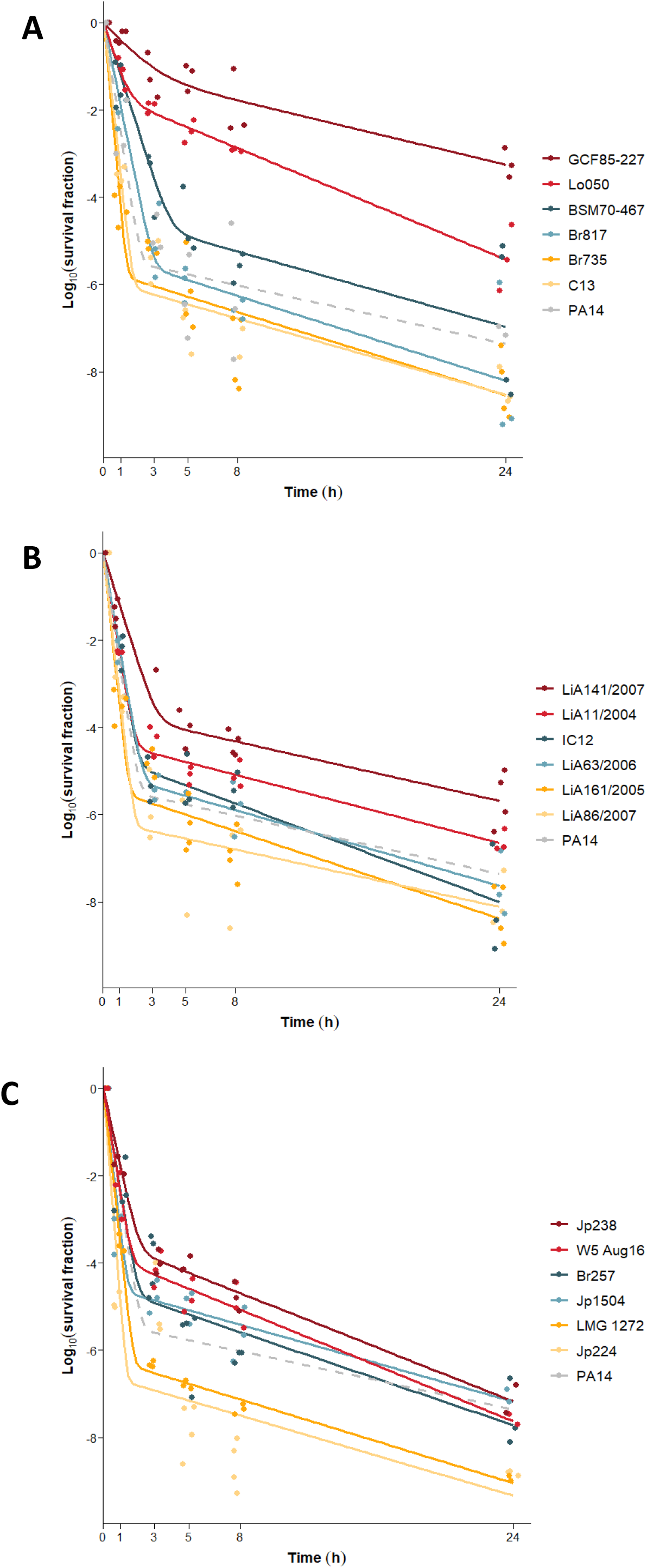
Time-kill curves of human (A), animal (B) and environmental (C) strains. Biphasic fittings of time-kill data in response to tobramycin treatment are shown in different colors according to their persistence phenotype. The colors red, blue and yellow indicate strains with, respectively, high, intermediate, and low persistence levels in vitro. Individual data points from at least three independent biological repeats are shown. The dotted line shows the killing kinetics of the lab strain PA14 which was included as a control in each experiment.

**Table 1:** The selection of natural *P. aeruginosa* isolates with their origin, susceptibility to tobramycin and survival group

|  |  |  |  | Origin |  |  |
| --- | --- | --- | --- | --- | --- | --- |
| Strain | MIC<br>tobramycin<br>(µg/ml) | *in vitro*<br>survival<br>phenotype | *in vivo*<br>survival<br>phenotype | Source |  | Country |
| C13 | 0.25 | Low | Low | Human | CF patient | Germany |
| BSM70-467 | 0.25 | Intermediate | High |  | CF sputum | Belgium |
| GCF85-227 | 0.125 | High | High |  | CF sputum | Belgium |
| Br817 | 0.25 | Intermediate | Intermediate |  | Wound | Belgium |
| Lo050 | 0.5 | High | High |  | Burn | United Kingdom |
| Br735 | 0.25 | Low | Intermediate |  | Burn wound | Belgium |
| LiA86/2007 | 0.25 | Low | Intermediate | Animal | Dog uterus | Portugal |
| IC12 | 0.25 | Intermediate | High |  | Dog | India |
| LiA63/2006 | 0.25 | Intermediate | Low |  | Kangaroo blood | Portugal |
| LiA141/2007 | 0.25 | High | High |  | Dog eye | Portugal |
| LiA161/2005 | 0.25 | Low | Low |  | Parrot eye | Portugal |
| LiA11/2004 | 0.25 | High | Intermediate |  | Cat nose | Portugal |
| Jp224 | 0.5 | Low | Low | Environmental | Sea water<br>(open ocean) | Japan |
| Jp1504 | 0.5 | Intermediate | High |  | River water | Japan |
| W5 Aug16 | 0.25 | High | Intermediate |  | River water | Belgium |
| LMG 1272 | 0.25 | Low | Low |  | Mushroom | United Kingdom |
| Br257 | 0.25 | Intermediate | Low |  | Plant rhizosphere | Belgium |
| Jp238 | 0.25 | High | Intermediate |  | Sea water (coastal) | Japan |

### *In vivo* survival of natural *P. aeruginosa* isolates

The representative strain collection was used to study the survival upon antibiotic exposure in the optimized murine model that mimics *P. aeruginosa* lung infections. In a first set of experiments, the optimized challenge dose of 5×10^5^ CFU/mouse was tested for all 18 isolates. All mice survived 26.5 h after infection with this dose (Figure S3A) and dissemination to liver and spleen was limited (Figure S3B & S3C). This challenge dose could thus be used for subsequent treatment experiments. In these experiments, mice were treated at 24 h p.i. with tobramycin (120 mg/kg mouse) via nasal droplets and sacrificed 2.5 h later. Euthanasia of untreated mice (control group) was performed at 26.5 h p.i. to make the infection process comparable to the one of treated mice (treated group) (Figure 1C). Mice treated with tobramycin showed a significant reduction of bacteria in the lungs compared to the control group with an average reduction across all infected isolates of 1 log (p = 0.02; data not shown). To be able to compare *in vitro* survival and *in vivo* survival, a similar quantitative measure was computed. The *in vitro* persister fraction is defined as the ratio of the number of surviving bacteria after treatment and the number of viable bacteria before treatment. In this study, the *in vivo* survival fraction in one mice is defined as the number of cells after treatment in the left lung of that mice divided by the average number of cells in the left lung of all untreated mice at 26.5 h p.i.:

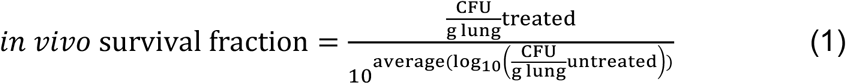

Similar analyses as for the *in vitro* survival fractions were performed on these calculated *in vivo* survival fractions. The observed *in vivo* survival fractions are variable among the different isolates (p < 0.0001) (Figure 4). However, in contrast to *in vitro* survival, there is no significant effect of the strain’s origin on the *in vivo* survival (p = 0.23). The isolates were again categorized into three groups based on their *in vivo* survival: low, intermediate and high (Table 1). The average survival fraction between these three groups is significantly different (p < 0.0001) (Figure S2B).

**Figure 4:**
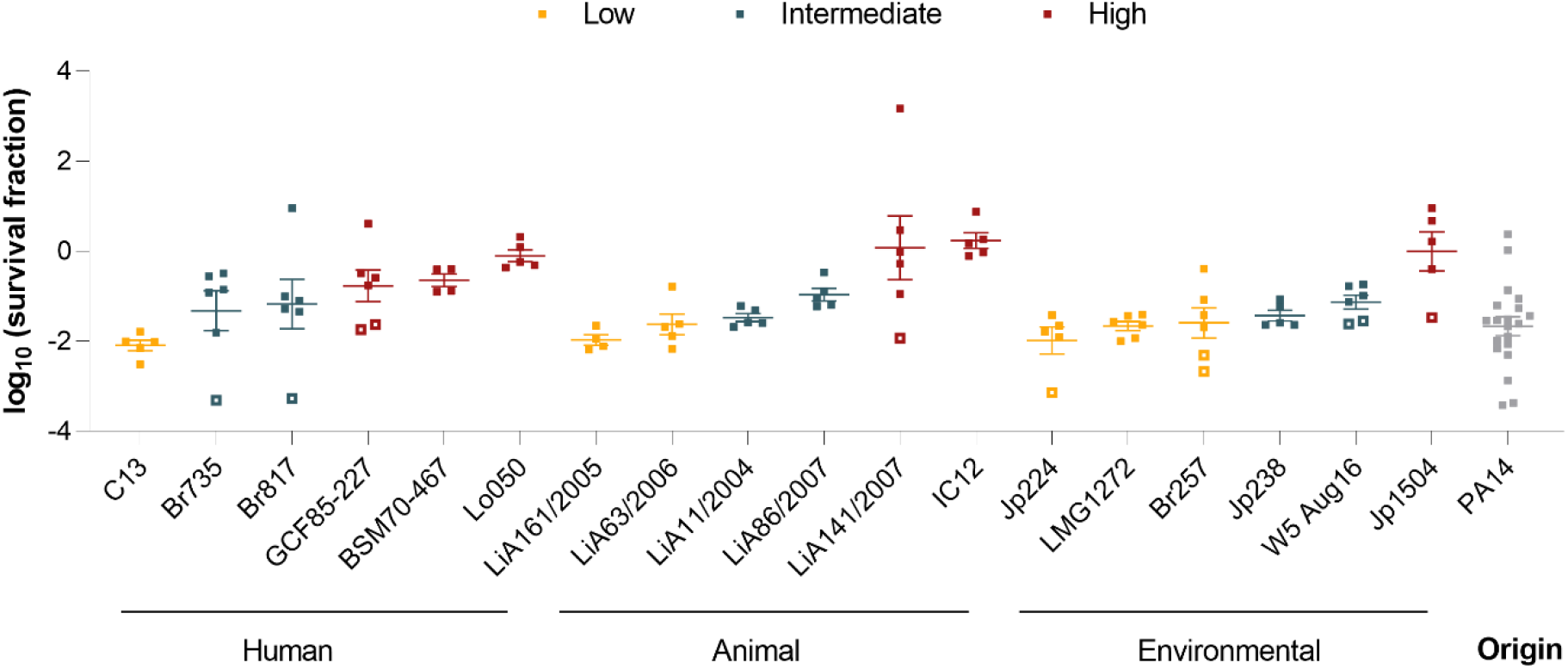
In vivo survival of P. aeruginosa isolates. Strains are sorted according to their origin and colored according to their in vivo survival phenotype with yellow, blue, and red representing, respectively, low, intermediate and high survival levels. Each square represents the survival fraction of individual mice observed in the murine lung infection model, calculated according to (1). Open squares indicate treated samples below the detection limit. Error bars represent the standard error of the mean.

To assess possible changes in host immunological responses upon antibiotic treatment and their impact on the observed bacterial survival levels, the expression of a selection of cytokines and chemokines in the lungs was examined via RT-qPCR (Figure S4). The expression levels of eight cytokines and chemokines was measured: interleukin-1β (IL-1β), tumour necrosis factor-α (TNF-α), IL-6, IL-10, keratinocyte chemoattractant (KC), macrophage inflammatory protein 2 (MIP-2), interferon-gamma-inducible protein 10 (IP-10) and monocyte chemoattractant protein 1 (MCP-1). More specifically, the gene expression before and after treatment in response to a selection of the *P. aeruginosa* isolates with a diverse *in vivo* survival level was examined: LMG 1272, LiA63/2006 and Br257 (low survival level); Jp238 and Br817 (intermediate survival level); Lo050 and LiA141/2007 (high survival level) with the lab strain PA14 included as control. In infected mice, the expression of IL-1β and KC was most upregulated. The effect of the lung infection seemed to be more moderate for the other cytokines (Figure S4). Tobramycin treatment did not significantly change the expression of any of the cytokines tested, except for IP-10 (p = 0.02). The level of this pro-inflammatory chemokine was significantly increased in treated mice infected with strain Br817 (p = 0.01) and LiA141/2007 (p < 0.0001). We also studied whether there was a difference in immune response between strains belonging to a different survival group (Figure S5). Mice challenged with high-survival strains demonstrated higher expression levels of KC and MCP-1 compared to the strains belonging to the low and intermediate survival group (p < 0.0001 for KC and MCP-1). In contrast, the expression of the chemokine MIP-2 was highest in the low survival group (p=0.04). For other cytokines, expression levels were similar in all survival groups. Overall, these mice experiments point to a diversity in bacterial survival upon tobramycin treatment between natural isolates, similar to what is observed *in vitro*. In the next step, survival *in vivo* is directly compared to survival *in vitro*.

### The antibiotic survival in lab and clinical settings is positively correlated

The 18 *P. aeruginosa* isolates can be ranked and categorized into three phenotype groups based on their *in vitro* and *in vivo* survival level: low, intermediate and high (Table 1). For 8 isolates, both the *in vivo* and *in vitro* phenotype groups are the same. The other 10 strains have switched from a low-persister or high-persister strain *in vitro* to an intermediate strain *in vivo*, and vice versa. This indicates that there are only small differences in categorization based on survival since strains did not switch phenotype groups or only changed to one group higher or lower. To further investigate the relationship between the *in vitro* and *in vivo* survival, a PGLS model was used to correct for non-independence between the isolates. First, the phylogenetic signal of the data set was quantitatively measured via Pagel’s λ and Blomberg’s κ. A λ or κ value of 1 illustrates a strong phylogenetic signal in the survival levels, a λ or κ value of 0 illustrates no phylogenetic signal [48]. Both metrics are close to zero which indicates a low, non-significant phylogenetic signal under Brownian model of evolution and thus closely related isolates do not have more comparable survival levels compared to distant isolates. However, it is difficult to estimate the phylogenetic signal for small datasets and low λ or κ does not necessarily mean that there is no phylogenetic signal [49–52]. We performed and compared two extreme models with fixed κ values and lambda = 1 and lambda = 0 (Figure 5, Table S2). The latter PGLS model is equivalent to an ordinary least squares (OLS) model. Both models revealed a strong dependency of the *in vivo* survival on the *in vitro* survival. The R^2^ of both models is 0.25 which means that 25% of the variation in *in vivo* survival can be explained by *in vitro* survival levels. Overall, our data suggest that the survival observed in laboratory settings is comparable to survival in conditions that mimic the clinical situation.

**Figure 5:**
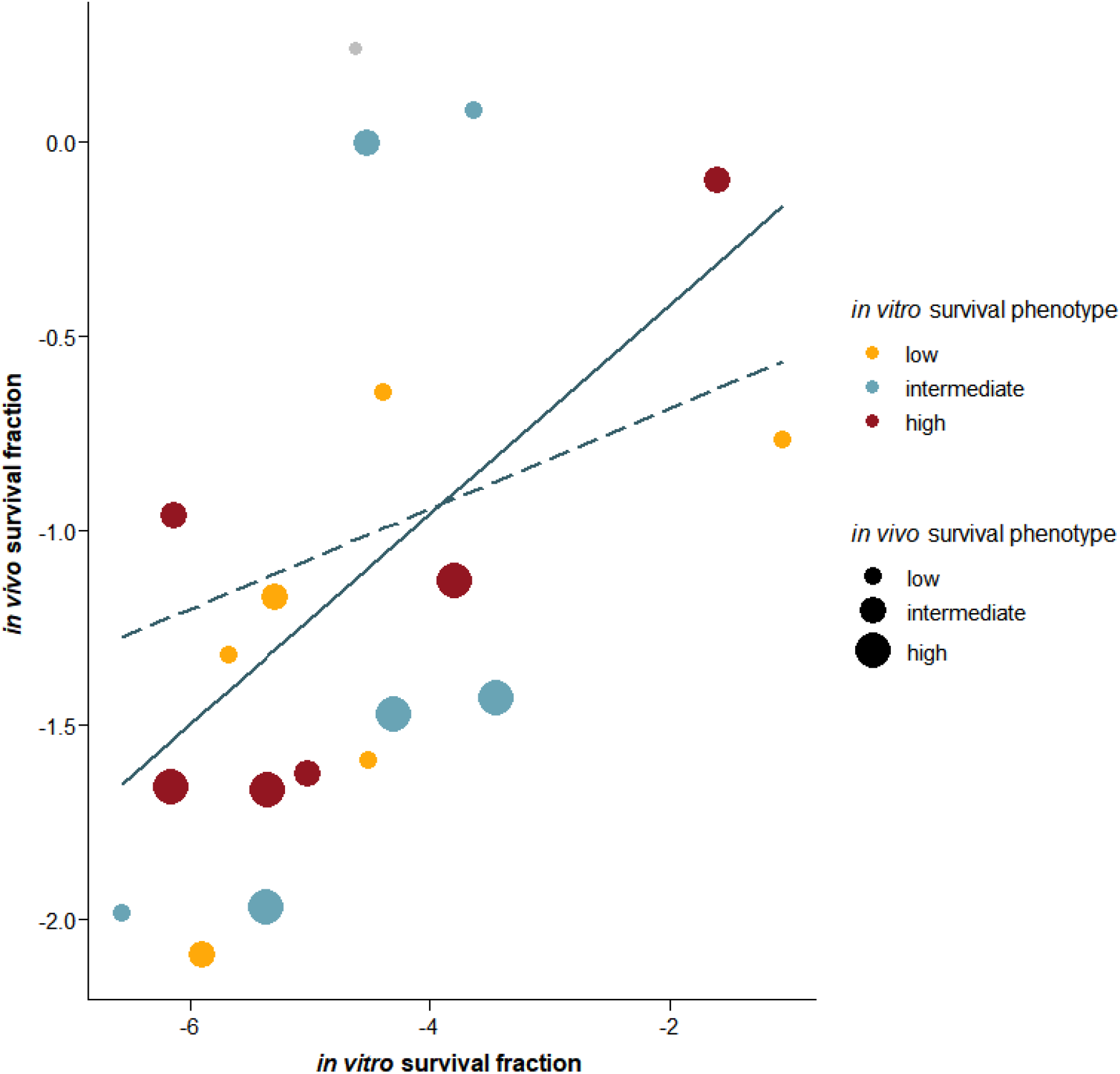
The in vitro survival is predictive for the in vivo survival. The mean survival fraction in both conditions is shown for all tested P. aeruginosa isolates. The 18 natural isolates are colored according to their phenotype in vitro, the size of the dots indicate their phenotype in vivo. The lab strain PA14 is represented by the grey dot. The solid line shows the OLS regression line and the dashed line the PGLS regression line.

## Discussion

Since its discovery in the 1940s by Hobby [53] and Bigger [54], persistence has gained great interest in the past years due to its role in relapsing infections [10] and resistance development [12–15]. This eventually led to the publication of a consensus statement in which a clear definition of persistence and general guidelines to measure persistence are provided [9]. One of the hallmarks to study persistence *in vitro* is the time-kill assay in which bacterial survival upon exposure to high antibiotic concentrations is followed over time. However, to date, it is not clear whether the results of these time-kill assays also resemble the antibiotic tolerance observed in patients. In this paper, we have developed a murine lung infection which allows us to study the *in vivo* tolerance of natural *P. aeruginosa* strains. Previous *in vivo* studies in persistence research have mainly focused on lab strains and persistence mutants with higher or lower persistence levels, but natural isolates have, to our knowledge, never been studied in *in vivo* persistence research. We found that *in vitro* survival can partly predict the survival in an *in vivo* setting and thus validates the use of time-kill assays to quantify persistence.

In the *in vitro* experiments, all *P. aeruginosa* isolates across all origins showed a rapid decrease of the majority of the cells with only a small fraction of cells surviving a high antibiotic treatment. We observed a large variation in persistence levels in natural isolates, similar as reported previously [15, 55–58]. Remarkably, the number of surviving cells was higher in clinical compared to environmental strains. Variation in persister formation among isolates may be explained by their evolutionary history [8]. The history of exposure to antibiotics or other environmental stresses of the examined isolates is unknown. Nevertheless, antibiotics, a known inducer of persistence, in natural environments is in most cases present in lower concentrations than in patients, even lower than the MIC [59]. It could be argued that strains isolated from clinical sources, such as CF lungs, experience higher selection pressure in comparison to strains isolated from non-clinical sources due to a prolonged exposure to high antibiotic doses. A similar effect clearly underlies the high-persistence of late isolates from long-term treated CF patients [6, 7]. Moreover, isolates from chronic *P. aeruginosa* infections show increased tolerance compared to isolates from acute infections [60].

We established a lung infection model in mice to study the survival of multiple *P. aeruginosa* isolates. Although there are some important anatomical and physiological differences between murine lungs and human lungs, this chronic infection model with bacteria embedded in agar, agarose or seaweed alginate beads is widely used to study the *in vivo* pathogenesis of lung infections and test the efficacy of new antibacterial therapies [61]. In acute infection models, free-living *P. aeruginosa* cells are directly administered to the murine lungs causing rapid clearance of the cells or acute sepsis and eventually death [43]. By embedding the bacteria in beads, bacterial clearance is avoided and the infection can be maintained for several weeks because there is a continued mild induction of the host immune system. Furthermore, the chronic model shows similarities with human pathology of chronic CF infections including the histopathology and the increase in lung neutrophils and cytokines such as IL-1β, TNF-α, MIP-2, IL-6 and KC [41, 40, 44]. The seaweed alginate bead model demonstrates a more pronounced antibody response compared to the agar bead model which is also typical for chronic *P. aeruginosa* infections. Moreover, such alginate beads are often preferred because they mimic mucoid *P. aeruginosa* biofilms where alginate is a constituent of the extracellular matrix [40].

In patients, antibiotic-tolerant biofilm structures make chronic infections very challenging to completely eradicate. The ability of biofilms to tolerate antibiotics has been attributed to several mechanisms including subpopulations with differences in metabolic activity, the presence of persisters, differential gene expression and limited diffusion of antibiotics through the biofilm matrix [62, 63]. The latter mechanism seems to be especially relevant for aminoglycosides since these positively charged antibiotics bind to the negatively charged extracellular matrix of biofilms. We offer several arguments to demonstrate that tobramycin effectively penetrates the alginate beads in our murine model. Cao et al. (2015) demonstrated that tobramycin still penetrates large seaweed alginate beads *in vitro* and even accumulates in the alginate matrix. Also several hours after removal of tobramycin from the beads the bacteria inside the beads were still effectively killed which shows that the concentration of free unbound tobramycin is still high [64]. More detailed pharmacokinetic analyses revealed that the tobramycin concentration in the alginate matrix follows a power law as function of the external concentration, probably by non-specific binding to the matrix [65]. These calculations should be taken with caution as this power-law dependence is calculated with *in vitro* alginate beads. Recently, Christophersen et al. (2020) tested these findings in a chronic lung infection model in mice in which they followed the tobramycin concentration and bacterial killing in the alginate beads over time. This study confirmed the slow release of tobramycin from the alginate beads. Unbound tobramycin was detected several hours after tobramycin administration at low concentrations [66]. It needs to be mentioned that direct comparison with our experiments is not possible as tobramycin in these experiments was administered subcutaneously at a lower dose. In all abovementioned studies, the alginate density was also kept constant at 3% and larger alginate beads were tested with a diameter ranging from 4 to 5 mm. Our alginate beads were made of 1% alginate, and have an average diameter of 25 μm (data not shown) which is more comparable with the colony size observed in *in vivo* biofilms [67]. Based on these arguments, we expect less retardation of diffusion through the beads in our optimized model but more elaborate pharmacokinetic and pharmacodynamic analyses are needed to determine the effective antibiotic concentration inside the alginate beads. Also, tobramycin is not completely degraded during the course of the experiment as pharmacokinetic profiles of tobramycin in an acute infection model showed that the concentration in the murine lungs remains high for at least 24 h after internasal administration [68]. Together, we believe that the administration of tobramycin for 2.5 h at an extreme high dose was sufficient to target all sensitive *P. aeruginosa* cells in the lungs.

In our lung infection model, tobramycin treatment significantly reduced the number of bacteria in the lungs. For all treatment durations, we observed a bacterial killing of PA14 with 2 log compared to the untreated group. Complete eradication upon tobramycin treatment in the seaweed alginate model was not observed which is consistent with previous research [66, 68, 69]. The timing of treatment probably plays an important role as total clearance of bacteria in the lungs is observed when antibiotics were administered immediately after infection [68, 69]. In the natural isolates, a less pronounced, but significant, reduction in CFU counts after treatment was observed. Although we also see a significant difference in survival level between the isolates, this variation is less pronounced than *in vitro* assays. Other factors may play a role in the observed tolerance of bacteria in chronic infections. In biofilms, cells undergo physiological and genetic alterations to survive in their new environment which affects antibiotic tolerance as well [62, 63]. Different than in *in vitro*, an effect of the strain’s origin on its *in vivo* survival was not observed. Bragonzi et al. demonstrated that CF strains and environmental *P. aeruginosa* strains have the same capacity to establish a chronic infection in murine lungs [70], but the effect of antibiotic treatment on strains with a different origin has not been demonstrated before. The cytokine profile revealed that upon tobramycin treatment no change in inflammatory response was detected which was also described earlier for tobramycin treatments starting later after infection [68]. Remarkably, increased KC, IP-10 and MCP-1 levels before and after treatment were observed in high-survival strains. While this is beyond the scope of this paper, this could hint at a direct relationship between virulence, which is here reflected by changes in immunological responses, and antibiotic tolerance. A possible link between virulence and persistence has been proposed previously [71]. Indeed, the stringent response, quorum sensing and toxin-antitoxin modules play an important role in persistence and also regulate the expression of virulence factors. The relationship between virulence and antibiotic tolerance seems an interesting topic for further investigation. For example, investigating the *in vivo* expression levels of virulence factors, such as elastase, protease A and pyocyanin, in strains with a variable tolerance level would be more informative. It is important to note that the increase in cytokine expression levels in our experiments seems only moderate. In future experiments, the inflammation of uninfected mice should be considered as well to allow comparison with normal expression levels of cytokines.

Despite the distinct characteristics between *in vitro* and *in vivo* set-ups, we still observe a correlation between the survival levels in both systems. We are aware that the optimized model still has its limitations and that the observed survival level cannot be solely explained by antibiotic tolerance. Nevertheless, strains with an extreme persistence phenotype have an extreme tolerant phenotype *in vivo* as well. For the first time, antibiotic tolerance between an *in vitro* and *in vivo* set-up is directly compared. Only one previous study on *P. aeruginosa* has confirmed the *in vivo* tolerant phenotype of a Δ*relA spoT* mutant upon ofloxacin treatment [21]. Inactivation of both genes disrupts the stringent response and results in lower persistence levels compared to the wild type *in vitro*. Intraperitoneal infection of the mutant increases the mice survival with 40% compared to the wildtype after antibiotic treatment which confirms the role of the stringent response in antibiotic tolerance [21]. Other murine models in persistence research were focused on testing the efficacy of a new antipersister therapy (e.g. [72–75]), unravelling the role of host factors (e.g.[28, 30, 76]) or tracking the metabolic activity with reporters for bacterial growth (e.g. [25, 28, 31]). The model described in this study allows us to both evaluate new drugs and study tolerance in a clinically relevant animal model.

## Materials and Methods

### Bacterial strains and culture conditions

Experiments were performed with natural *P. aeruginosa* strains isolated from clinical, environmental, or animal sources (Table 1). The lab strain UCBPP-PA14 (PA14) was used for the optimization of the *in vivo* model and included in each subsequent experiment. For the *in vitro* experiments, bacterial cultures were grown at 37°C in Mueller-Hinton broth (MHB; BD Difco) with orbital shaking or on lysogeny broth (LB; VWR International) agar. For the preparation of the seaweed alginate beads, *P. aeruginosa* strains were grown at 37°C in tryptic soy broth (TSB; BD Difco) with orbital shaking (150 rpm).

### Production of *P. aeruginosa* embedded in seaweed alginate beads

The encapsulation of *P. aeruginosa* in seaweed alginate beads was performed as previously described with minor modifications [41, 77]. One colony of the *P. aeruginosa* strain to be tested was incubated for 18 h in 50 ml of TSB at 37°C. The overnight bacterial culture was then centrifuged at 4750 rpm for 10 minutes at 4°C and the pellet was resuspended in 5 ml TSB. 600 μL of this bacterial culture was suspended in 12 mL of sterile 1% alginate (FMC Biopolymer). The *P. aeruginosa-* alginate suspension was then transferred to a 20 ml syringe (BD Plastipak) fixated on a syringe pump (Prosense, Multi-Phaser) which was connected to the Encapsulation Unit Var J30 (Nisco). The suspension passed through the nozzle at a flow rate of 12 ml/h into a gelleting bath. This gelleting bath contained 0.1 M Tris-HCl buffer and 0.1 M CaCl2 (pH 7, Sigma) and was continuously mixed by magnetic stirring to avoid merging of the alginate beads. After 1 h of stirring in the gelleting bath, the alginate beads were washed twice with 0.9% NaCl containing 0.1 M CaCl2 and suspended in this NaCl buffer. To determine the number of bacteria inside the beads, the alginate beads were dissolved using 0.5 M citrate buffer, serially diluted, plated on LB agar plates, and incubated for 24 hours. The concentration of bacteria in the beads was approximately 10^7^-10^8^ CFU/ml. The alginate beads were visualized by phase contrast microscopy using Nikon Eclipse Ti-E inverted microscope equipped with an Qi2 CMOS camera.

### Determination of the minimum inhibitory concentration

The minimum inhibitory concentration (MIC) of an antibiotic is the lowest concentration at which the growth of the bacterial strain is inhibited and indicates the susceptibility of a strain to this antibiotic. To determine the MIC of tobramycin (TCI) of the isolates, the microbroth dilution in 96-well microtiter plates or the agar dilution method was performed [78]. For the microbroth dilution method, an overnight culture was diluted in MHB to approximately 5 x 10^5^ cells/ml and incubated in a twofold dilution series of tobramycin at 37°C for 24 h. The MIC was determined by measuring the optical density at 595 nm with a Synergy Mx Microplate Reader (BioTek). For the agar dilution method, Mueller Hinton agar dilution plates were prepared with a concentration ranging from 0.0313 μg/ml to 64 μg/ml with antibiotic-free plates as a positive control. An overnight culture was adjusted to a cell density of approximately 10^6^ CFU/ml and subsequently, 10 μl of the suspension was spotted on the agar dilution plates resulting in a final inoculum of 10^4^ CFUs per spot. Plates were incubated at 37°C for 20 to 24 h after which the MIC was determined by visual inspection of colony formation. The MIC of each strain was determined based on at least two independent biological repeats.

### *In vitro* killing assays

Strains were inoculated in 500 μl MHB in deep-well plates (U-bottom) and incubated at 37°C for 24 h on a Titramax 1000 shaker (Heideloph) at 1200 rpm. These cultures were then 1:100 diluted in fresh MHB medium and incubated under the same growth conditions. After 16 h, the cultures were 1:2 diluted in fresh medium after which an aliquot of 200 μl was treated with tobramycin (100x MIC, concentration adjusted per strain). After 1, 3, 5, 8 and 24 h of incubation, the bacterial cells were washed twice with 10 mM MgSO_4_ to remove remaining antibiotics. The number of CFUs before and after antibiotic treatment were determined by plating on LB agar. The plates with untreated cultures were counted after 24 h of incubation at 37°C, plates with treated cultures after 48 h.

### *In vivo* killing assays

All animal experiments were authorized and approved by the Ethical Committee of the University of Antwerp (approval number 2018–91). 252 female BALB/c mice of 12 weeks old (Janvier Labs) were managed in accordance to the guidelines provided by the European Directive for Laboratory Animal Care (Directive 2010/63/EU of the European Parliament). Mice were briefly sedated with isoflurane (Halocarbon) and then intratracheally infected with *P. aeruginosa* embedded in seaweed alginate beads. The optimized challenge dose was 5×10^5^ CFU/mouse. For this procedure, mice were briefly sedated with isoflurane and 40 μl of the alginate solution was administered above the vocal cords. After 24 h of infection, a group of mice received one dose of approximately 120 mg/kg body weight tobramycin via intranasal application. Mice were held in a supine position and 20 μl of tobramycin solution was administered in each nostril. Upon regain of consciousness, mice had unlimited access to feed and water. Body weight was monitored daily since mice that lose ≥ 20% bodyweight are considered in morbid status and must be euthanized. Both untreated and treated mice were sacrificed at 24 h, 26.5 h, 29 h or 31.5 h p.i. by cervical dislocation. The left lung, left lateral lobe of the liver, and the entire spleen were harvested and homogenized in 1 ml PBS using TissueRuptor (Qiagen). These homogenates were serially diluted and plated on agar plates to determine the number of colonized bacteria in each organ. The bacterial burden was expressed per gram of organ.

### Phylogenetic analysis

Raw sequencing data of strains Br735, LiA63/2006, LiA161/2005, Jp224, W5 Aug16 and Jp238 are available on the International Pseudomonas Consortium Database (IPCD) and at the National Center of Biotechnology Information (NCBI). The whole-genome sequencing of the other strains was performed on an Ilumina MiniSeq (Laboratory of Gene Technology, KU Leuven, Belgium) or NextSeq 500 (Genomics Core, UZ Leuven, Belgium). Trimmomatic v0.39 was used for read trimming and filtering including adapter removal, trimming based on quality (LEADING:3; TRAILING:30; SLIDINGWINDOW:4:20) and removal of reads shorter than 36 bases [79]. The quality of the reads before and after trimming and filtering was assessed using FastQC v0.11.8. The genomes were assembled *de novo* from the trimmed paired-end reads using SPAdes assembler v3.14.1 [80] and the quality of the draft genome was assessed with QUAST v5.0.2 [81]. The obtained genome sequences were annotated with Prokka v1.14.6 using the GenBank compliance option which removes contigs shorter than 200 bases [82]. Core genes, which are present in 99% of the strains, were extracted and analyzed using Roary v3.13.0 with default settings [83]. The resulting core-gene alignment was used to construct a phylogenetic tree with IQ-TREE v2.1.4 with automatic model selection [84, 85] and ultrafast bootstrapping (UFBoot) [86]. Molecular clock rate estimates were generated with TreeTime and used for the construction of an ultrametric tree [87]. iTol was used for tree visualization [88].

### Characterization of cytokine respones by quantitative PCR

After sacrificing the mice, the inferior lobe of the right lung was collected in RNAlater (Invitrogen). Total RNA was extracted using TRIzol reagent according to the manufacturer’s protocol. In short, samples were thawed and homogenized in TRIzol reagent (Invitrogen) using a Qiagen TissueRuptor. After addition of chloroform, samples were centrifuged and the transparent layer was transferred to a new centrifuge tube to which isopropanol was added for RNA precipitation. The samples were then washed with 75% ethanol and air-dried for 5-10 minutes. The pellet was resuspended in nuclease free water and the concentration was measured using a NanoDrop 2000 spectrophotometer (Thermo Scientific). The extracted RNA was treated with ezDNase Enzyme (Invitrogen) according to the manufacturer’s protocol to remove any remaining genomic DNA. For the cDNA synthesis, SuperScript IV first-strand synthesis system (Invitrogen) was used following the manufacturer’s protocol. Targets were amplified using oligo(dT)-primers to select for mRNA specifically by binding to the poly(A)-tail. Both reverse transcriptase (RT) and no-RT control reactions were performed as a control for the presence of remaining genomic DNA during the subsequent PCR reaction. Quantitative PCR (qPCR) was performed using specific primer-probe pairs in 3 multiplex panels. These panels consisted of Th1, Th2 and chemokine specific pairs. Experiments were performed on a LightCycler 96 (Roche) using the SensiFAST Probe No-ROX kit (Bioline). All data was analyzed with the LightCycler 96-software and normalized by comparing them to the expression of two housekeeping genes (GAPDH and β-actin).

### Quantification of tobramycin concentrations in the lungs

Tobramycin concentrations in the murine lungs were determined via the agar well diffusion method with *Bacillus subtilis* ATCC 6051 as the indicator strain [89]. A standardized inoculum of 10^8^ cells/ml was diluted 1:200 in 1% MHB agar before pouring the agar plates. Holes with a diameter of 8 mm were punched in the solidified agar and 100 μl of standard tobramycin solution or lung homogenate were introduced into the wells. After overnight incubation at 37 °C, the diameters of the inhibition zones were measured with a caliper. A standard curve constructed with standard tobramycin solutions, ranging from 0.0625 μg/ml to 32 μg/ml, was used to determine the tobramycin concentration of the lung homogenates. This assay was carried out in triplicate with lungs infected with PA14 from three different mice experiments, each consisting of two technical repeats.

### Statistical analysis

All statistical tests were performed on log10 transformed data using GraphPad Prism v9.0.0 or R v.4.1.2. When the lowest dilution yielded a CFU count of 0, data points were assigned with a value of half the detection limit. The Shapiro-Wilk test was used to test for normality of the data. Significant changes were assessed through a one-way/two-way analysis of variance (ANOVA) if the variables were normally distributed. In case of a non-normal distribution, the Kruskal-Wallis test was used. Biphasic nonlinear mixed models were fitted to the time-kill data using the R package *nlme* based on the following equation:

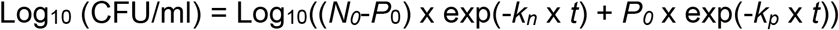

With *t* the treatment time (in hours), *k_n_* and *k_p_* the killing rate of normal and persister cells (per hour), respectively, and *N_0_* and *P_0_* the number of cells (in CFU) at *t* = 0. The biphasic models were compared with uniphasic nonlinear mixed models (with *P_0_* = 0) using the Akaike Information Criterion (AIC). The model with the lowest AIC score indicates a better fit. Pylogenetic generalized least squares (PGLS) analyses using the R packages nlme/ape, caper, phytools and picante were performed to account for phylogeny when testing the association between survival levels. The constructed ultrametric phylogenetic tree and survival levels were used as input. The determination coefficient (R^2^) of the PGLS model was calculated as the R^2^_pred_ in the R package ‘rr2’ [90].

## Data availability

Raw sequencing data of strain Br735 is available at NCBI under Bioproject no. PRJNA297679 (SRR2939473). Sequencing data of strains LiA63/2006, LiA161/2005, Jp224, W5 Aug16 and Jp238 are available on IPCD (https://ipcd.ibis.ulaval.ca/) and at NCBI under Bioproject no. PRJNA325248. Sequencing data of the other strains is available at NCBI under Bioproject no. PRJNA XXXXXX.

## Acknowledgments

The authors would like to thank Niels Høiby and Claus Moser for hosting J.A. in their laboratory in the department of clinical microbiology at the Rigshospitalet, Copenhagen for training regarding the mouse model, and Lars Christophersen for his technical support during this training. We also thank Pierre Cornelis for kindly providing the *P. aeruginosa* UCBPP-PA14 strain, and Jean-Paul Pirnay and Daniel De Vos (Laboratory for Molecular and Cellular Technology, Queen Astrid Military Hospital) [91] and Françoise Van Bambeke (Pharmacologie Cellulaire et Moléculaire, UCLouvain) for kindly providing the natural *P. aeruginosa* isolates. We also thank Gabriel Perron and Roger C. Levesque for providing us the raw sequencing data of an isolate subset. The phylogenetic analyses were performed using the infrastructure of the Flemish Supercomputer Center (VSC).

J.A. was funded by the research project PRINTAID, the EU Framework Programme for Research and Innovation within Horizon 2020 - Marie Sklodowska-Curie Innovative Training Networks under grant agreement No. 722467. LV and BVdB received a scholarship from Research Foundation Flanders (FWO; 12O1917N, 12O1922N,1S86721N). The work was supported by grants from FWO (G055517N, 1528318N, 1513120N), KU Leuven (C16/17/006) and Flanders Institute for Biotechnology (VIB).

## Supplementary information

**Table S1:** Estimates of *in vitro* survival fraction based on biphasic kill fits

| Strain | Estimate of<br>$\log_{10}(\text{survival fraction})$ | 95% confidence intervals |
| --- | --- | --- |
| Br257 | -4.51 | [-4.01,5.02] |
| Br735 | -5.68 | [-5.18,-6.18] |
| Br817 | -5.29 | [-4.60,-5.97] |
| BSM70-467 | -4.38 | [-3.66,-5.11] |
| C13 | -5.91 | [-5.34,-6.47] |
| GCF85-227 | -1.05 | [-0.07,-2.03] |
| IC12 | -4.62 | [-4.07,-5.16] |
| Jp1504 | -4.53 | [-3.95,-5.10] |
| Jp224 | -6.57 | [-6.05,-7.10] |
| Jp238 | -3.45 | [-2.86,-4.03] |
| LiA11/2004 | -4.30 | [-3.75,-4.85] |
| LiA141/2007 | -3.64 | [-3.01,-4.26] |
| LiA161/2005 | -5.37 | [-4.87,-5.87] |
| LiA63/2006 | -5.02 | [-4.38,-5.67] |
| LiA86/2007 | -6.14 | [-5.56,-6.71] |
| LMG 1272 | -6.16 | [-5.59,-6.74] |
| Lo050 | -1.60 | [-1.04,-1.60] |
| W5 Aug16 | -3.79 | [-3.21,-4.37] |
| PA14 | -5.35 | [-4.77,-5.93] |

**Figure S1:**
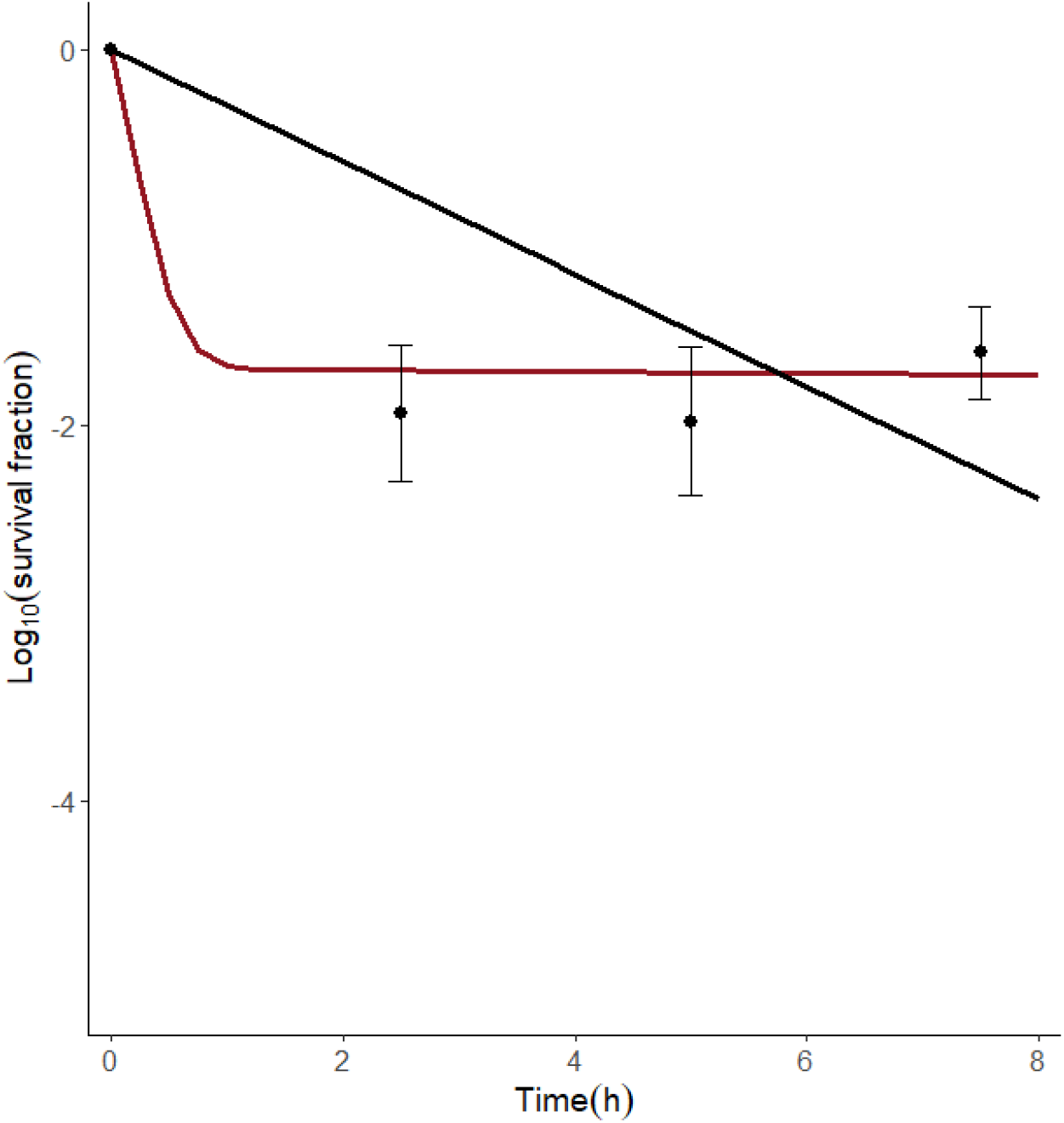
Time-kill curve of the PA14 murine lung population after tobramycin treatment. Both a biphasic exponential curve (in red) and an uniphasic exponential curve (in black) were fit to the time-kill data of Figure 1. Based on the Akaike Information Criterion (AIC) values, the biphasic model is superior over the uniphasic model to describe the observed killing dynamics.

**Figure S2:**
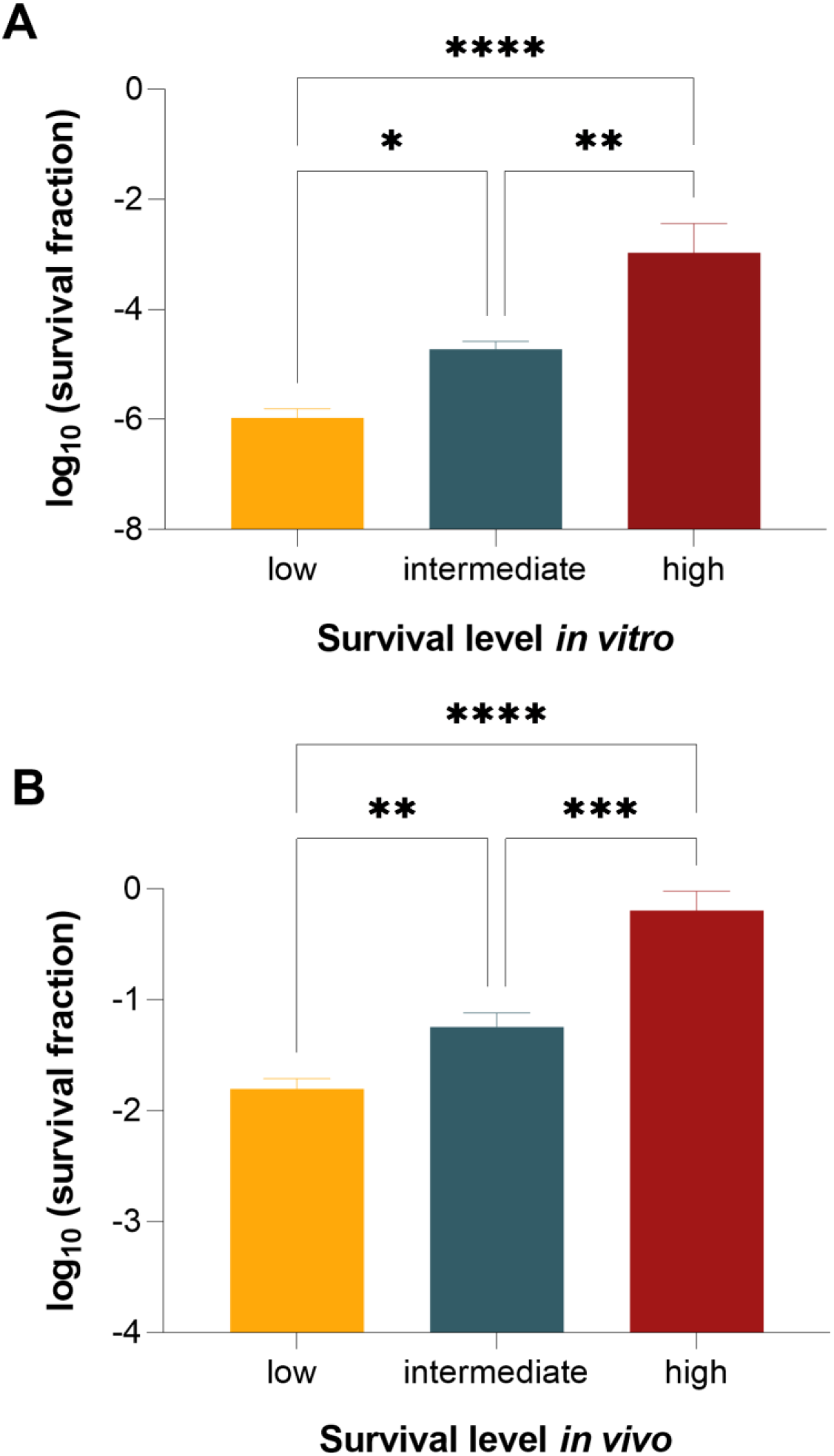
The survival fraction between the three survival groups is significantly different. Bars are coloured according to their survival level in vitro (A) and in vivo (B). Error bars show the standard error of the mean. Statistical differences between the in vitro groups are determined with one-way ANOVA followed by Tukey’s post hoc test for multiple comparisons. Statistical differences between the in vivo groups are determined via Kruskal-Wallis test followed by Dunn’s post hoc test for multiple comparisons. *, P < 0.05; **, P < 0.01; ****, P < 0.0001.

**Figure S3:**
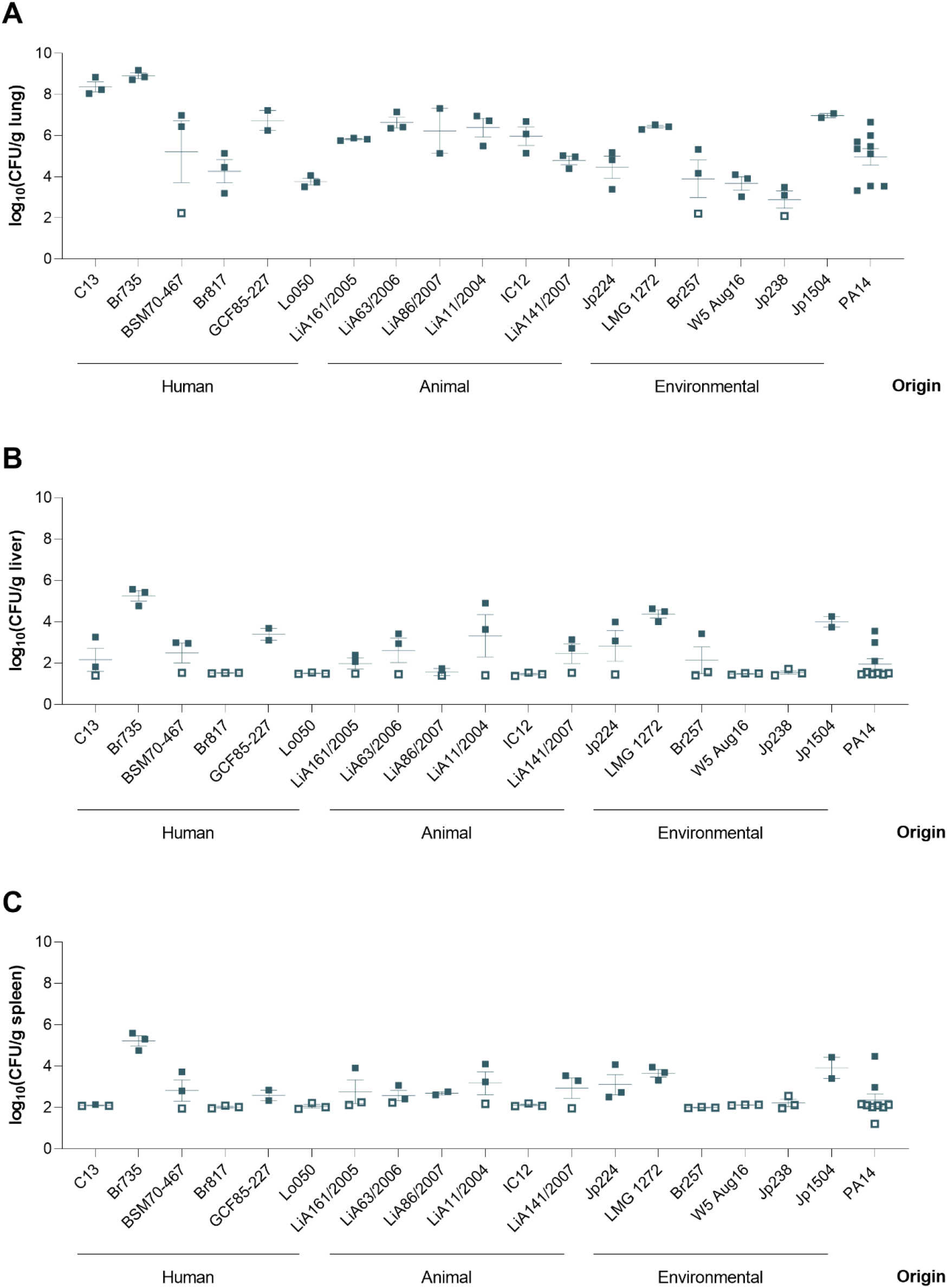
Bacterial load in the left lung (A), liver (B) and spleen (C) at 26.5 h p.i. of untreated mice. Open squares indicate repeats below the detection limit of which half of the detection limit divided by the organ weight is shown.

**Figure S4:**
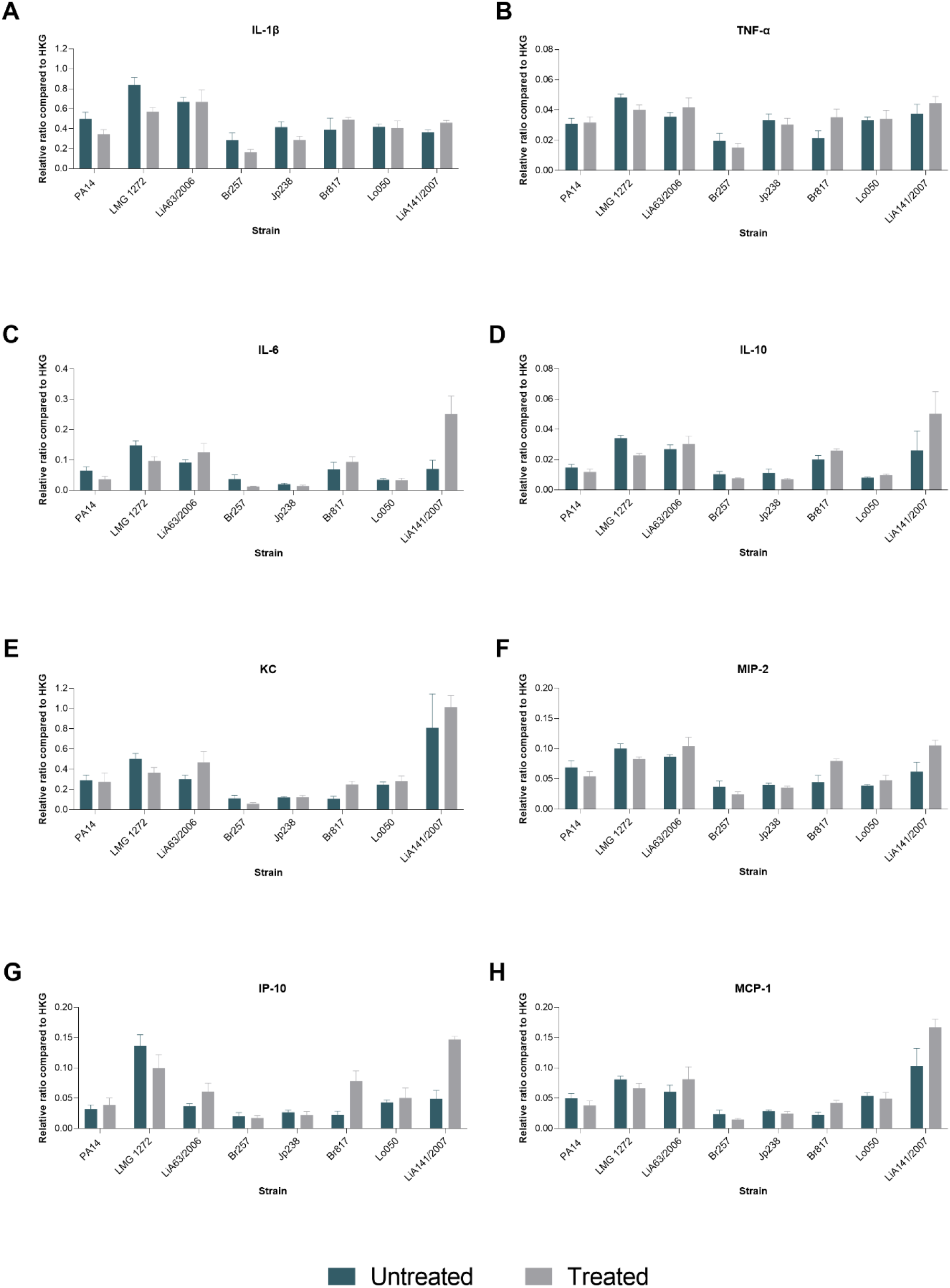
RT-qPCR analyses on a selection of isolates to determine the cytokine and chemokine expression levels. Quantitative PCRs were performed on the right lungs of mice (5-6 mice per group) infected with different P. aeruginosa isolates. The mice were sacrificed 26.5 h post infection with or without tobramycin treatment as indicated by the colours. Strains are sorted according to their in vivo survival, PA14 has the lowest survival and LiA141/2007 the highest survival. Expression levels are expressed relative to the expression of two housekeeping genes (GAPDH and β-actin). Except for IP-10, there is no significant difference in inflammatory response upon antibiotic treatment. Statistical analyses were performed per cytokine/chemokine with two-way ANOVA. Error bars represent the standard error of the mean.

**Figure S5:**
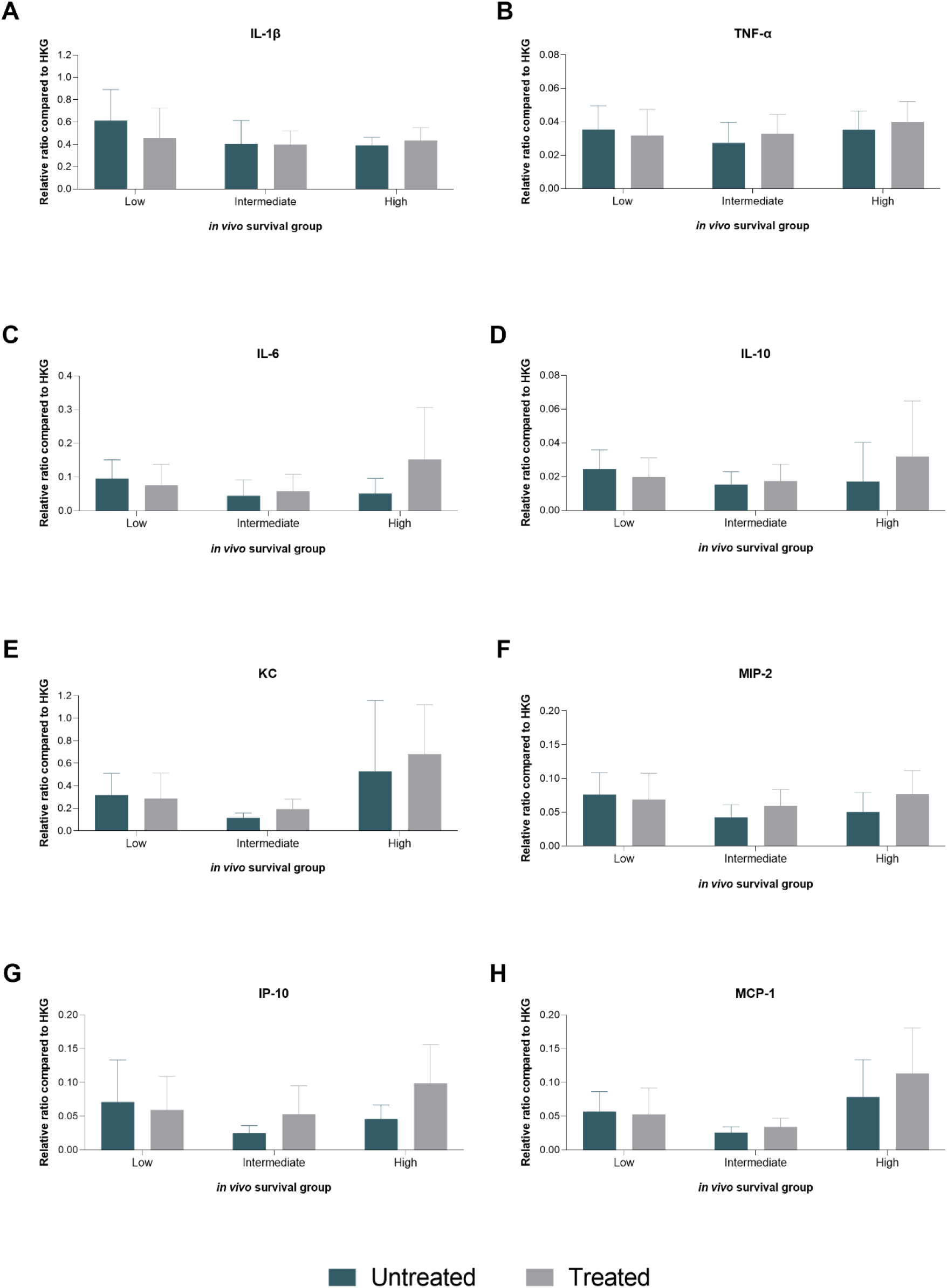
Cytokine and chemokine expression levels following P. aeruginosa infection classified per survival group. The same expression levels as Figure S4 are presented here, but are now shown per in vivo survival group instead of per strain. Strains are sorted according to their in vivo survival group, with the low survival group representing strains with the lowest survival fraction in vivo (LMG 1272, LiA63/2006 and Br257) and the high survival group representing strains with high survival fractions in vivo (Lo050 and LiA141/2007). The strains Jp238 and Br817 belong to the intermediate survival group. The lab strain PA14 was not included here for data analysis. The expression levels of KC and MCP-1 are significantly higher in the high survival group compared to the two other survival groups. The expression level of MIP-2 was highest in the low survival group. No significant differences between survival groups are observed for the other cytokines/chemokines. Statistical analyses were performed per cytokine/chemokine with two-way ANOVA and Šídák’s post hoc test for multiple comparisons. Error bars show the standard deviation.

**Table S2:** Comparison of the OLS model and PGLS model. SE = standard error, CI = confidence interval, AIC = Akaike Information Criterion

| regression<br>model | $\lambda/k$ | intercept | SE | 95% CI | p-value | slope | SE | 95% CI | p-value | R<sup>2</sup> | AIC |
| --- | --- | --- | --- | --- | --- | --- | --- | --- | --- | --- | --- |
| OLS | 0 | 0.1168 | 0.4988 | [-0.9355,1.1691] | 0.8177 | 0.2691 | 0.1037 | [0.0504,0.4879] | 0.01880 | 0.2417 | 40.813 |
| PGLS | 1 | -0.4302 | 1.9667 | [-4.5673,3.7068] | 0.8294 | 0.1288 | 0.0465 | [0.0307,0.2269] | 0.01309 | 0.2490 | 86.118 |

